# Elastin-derived peptides suppress CCL20 expression and block ILC2 recruitment during lung inflammation

**DOI:** 10.64898/2026.06.18.733133

**Authors:** Sarah Lahire, Caroline Fichel, Lisa Prince, Jeanne-Marie Perotin, Gaétan Deslée, Sébastien Le Jan, Stéphane Potteaux, Richard Le Naour, Arnaud Pommier

## Abstract

Elastin degradation during chronic lung inflammation generates elastin peptides (EPs) with immunomodulatory properties. Because elastin is abundant in the lung, its breakdown in diseases such as chronic obstructive pulmonary disease (COPD) and asthma produces high EPs levels that may influence local immune responses. Here, we investigated the impact of EPs on group 2 innate lymphoid cells (ILC2) using mouse models of EP-induced emphysema and house dust mite (HDM)–induced asthma. EPs instillation reduced lung ILC2 numbers without affecting Th2 cells. In patients with COPD, we observed decreased CCL20 expression in lung immune cells and an inverse correlation between serum CCL20 levels and clinical indicators of elevated EPs burden. We also showed that EPs instillation during HDM-induced lung inflammation directly decreased CCL20 expression. These findings identify EPs as regulators of ILC2 trafficking through CCL20 downregulation, revealing a direct link between extracellular matrix (ECM) degradation and the chemokine networks orchestrating type 2 immunity.

**One Sentence Summary:** Elastin-derived peptides reshape type 2 immunity by blocking CCL20-driven ILC2 recruitment during lung inflammation.

## INTRODUCTION

Lung immunity is essential for maintaining respiratory function and preserving barrier integrity (*1*). Type 2 immune responses play a central role in epithelial repair and protection against helminths and viral infections, but their dysregulation drives allergic airway inflammation and chronic lung disease (*2*). Group 2 innate lymphoid cells (ILC2) are key orchestrators of these processes: they contribute to host defense and tissue repair (*3*), yet also promote pathological type 2 inflammation (*4*). At steady state, pulmonary ILC2 are seeded early in life, remain tissue-resident, and self-renew locally (*5*). During inflammation, however, additional ILC2 can be recruited from circulation or lymphoid tissues into the lung parenchyma (*6*, *7*), where they amplify type 2 immunity and contribute to both protective (*8*) and pathogenic responses (*9*). Thus, the ability of ILC2 to accumulate in the lung at the onset of inflammation is a critical determinant of the magnitude and outcome of type 2 immune responses.

During lung inflammation, extensive remodeling of the ECM generates bioactive fragments known as matrikines, which can shape immune responses and promote type 2 inflammation (*10*). Among these, elastin-derived peptides (EPs) have been shown to modulate the activity of multiple immune cell types, including monocytes (*11*), neutrophils (*12*), NK cells (*13*), and T cells (*14*). Elastin is particularly abundant in the lung, and its protease-mediated degradation by recruited myeloid cells contributes to tissue destruction and emphysema (*15*). As a result, EPs accumulate in both acute and chronic lung injury (*16*). Several studies have demonstrated that EPs influence innate (*17*, *18*) and adaptive (*19*, *20*) immune responses during lung inflammation; however, aside from our recent work showing increased circulating ILC2 in patients with COPD (*21*), the impact of EPs on ILCs behavior during lung inflammation remains poorly understood.

Our earlier work established that EPs shape inflammatory responses in the lung (*20*, *22*, *23*) and increase circulating ILC2s in patients with COPD (*21*), raising the possibility that they may also regulate ILCs behavior within the lung. Given their abundance in inflamed tissue and their capacity to modulate diverse immune populations, EPs could, in principle, influence several aspects of ILCs biology. Yet it remained unclear whether EPs affect ILCs activation, expansion, subset identity, or their ability to accumulate in the lung during inflammation.

To address this question, we systematically examined how EPs influence the cellular processes that govern ILC accumulation in the lung. We assessed whether EPs alter ILC subset composition and the ability of ILCs—particularly ILC2—to populate inflamed lung tissue. Because ILCs recruitment is tightly controlled by chemokine cues, we also investigated whether EPs reshape chemokine pathways involved in ILC2 positioning.

## RESULTS

### EPs selectively limit the accumulation of lung ILC2s during inflammation

To determine how EPs influence innate lymphoid cells, we instilled EPs or PBS intratracheally and analyzed lungs, mediastinal lymph nodes (mLNs), and peripheral lymph nodes (pLNs) 3, 7, 14, and 21 days later (Fig. S1–S2). Consistent with our previous work showing that EP instillation triggers lung inflammation and leads to emphysema (*20*, *22*), EP-treated mice displayed clear signs of inflammation including a marked accumulation of Gr1⁺ myeloid cells at day 7 (Fig. S3) and increased numbers of Tbet⁺ CD8⁺ and CD4⁺ T cells at days 7 and 14, as well as GATA3⁺ CD4⁺ T cells at days 14 and 21 (Fig. S4–S5). PBS instillation also induced a mild inflammatory response, reflected by a progressive rise in lung Gr1⁺ cells (Fig. S3), which was accompanied by a transient accumulation of lung ILC2s at days 3 and 7 (Fig. 1A). Strikingly, despite inducing inflammation of greater magnitude, EPs completely prevented this ILC2 accumulation (Fig. 1A). Instead, EP-treated mice showed increased ILC2 numbers in the mLNs (Fig. 1B). Other ILCs subpopulations showed no EP-dependent changes in lung or lymph nodes (Fig. S6). Together, these data demonstrate that EPs selectively limit the inflammatory accumulation of lung ILC2s without broadly suppressing lymphoid responses.

**Figure 1:**
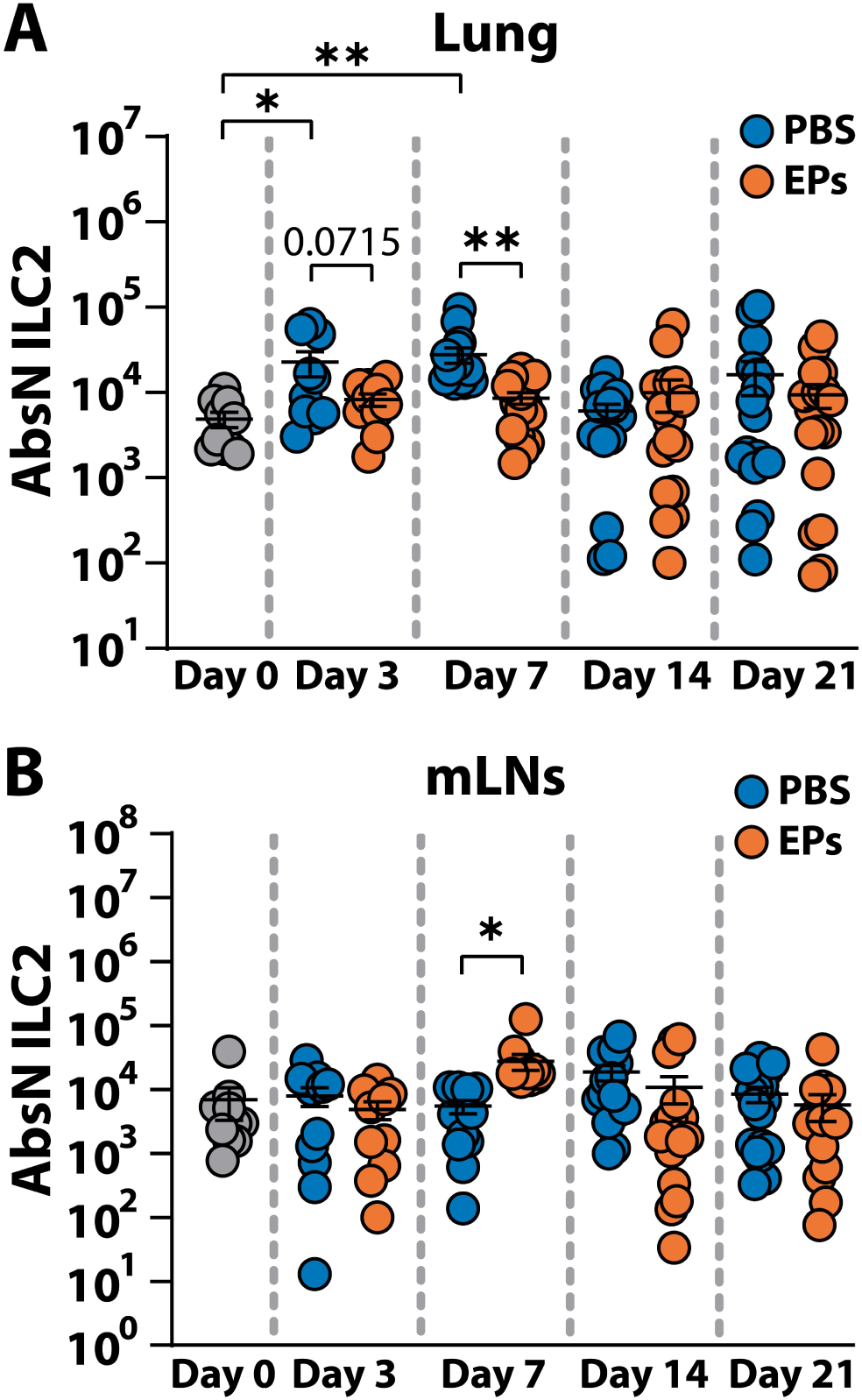
Lungs and mediastinal lymph nodes ILC2 numbers after EPs treatment. Mice were intratracheally instilled with 10 μg EPs in 50 μL in PBS or 50 μL of PBS alone. 0 (n=10), 3 (PBS n=10, EPs n=10), 7 (PBS n=16, EPs n=17), 14 (PBS n=15, EPs n=17) or 21 (PBS n=19, EPs n=18) days after, mice were exsanguinated by cardiac perfusion and lungs and lymph nodes were harvested to be analyzed by flow cytometry. Absolute number of **(A)** lung ILC2 and **(B)** mediastinal lymph nodes ILC2. Data are presented as scatter plots with mean ± SEM. Differences between two groups were evaluated by using the Wilcoxon-Mann–Whitney test. P values <0.05 were considered significant.

### EPs prevent ILC2 trafficking to the lung

We next investigated why ILC2s fail to accumulate in the lung during EP-induced inflammation. We considered three non-exclusive possibilities: impaired proliferation, conversion into other ILC subsets, or defective trafficking. To test proliferation, we adoptively transferred CFSE-labeled CD45.1⁺ ILCs into CD45.2 recipients 3 days before EPs treatment, and assessed CFSE dilution 7 or 14 days after PBS or EP instillation. Donor ILC2s proliferated similarly in both conditions in the lung and mLNs, indicating that EPs do not impair ILC2 expansion (Fig. S7). Furthermore, the quantification of GATA3⁺Tbet⁺ and GATA3⁺RORγt⁺ double-positive ILCs 7 days after EPs treatment revealed no EP-dependent differences in subset conversion (Fig. S8).

We therefore examined whether EPs interfere with ILC2 trafficking. CD45.1⁺ enriched ILCs were adoptively transferred 3 days after PBS or EPs instillation, so that they are already in an inflammatory environment, and donor-derived ILC2s were quantified 7 or 14 days later (Fig. 2A and Fig. S9). EP treatment almost completely abrogated ILC2 recruitment to the lung at day 7 and markedly reduced it at day 14, while donor ILC2 numbers in the mLNs remained unchanged (Fig. 2B–C). These results demonstrate that EPs block the entry of circulating ILC2s into the inflamed lung.

**Figure 2:**
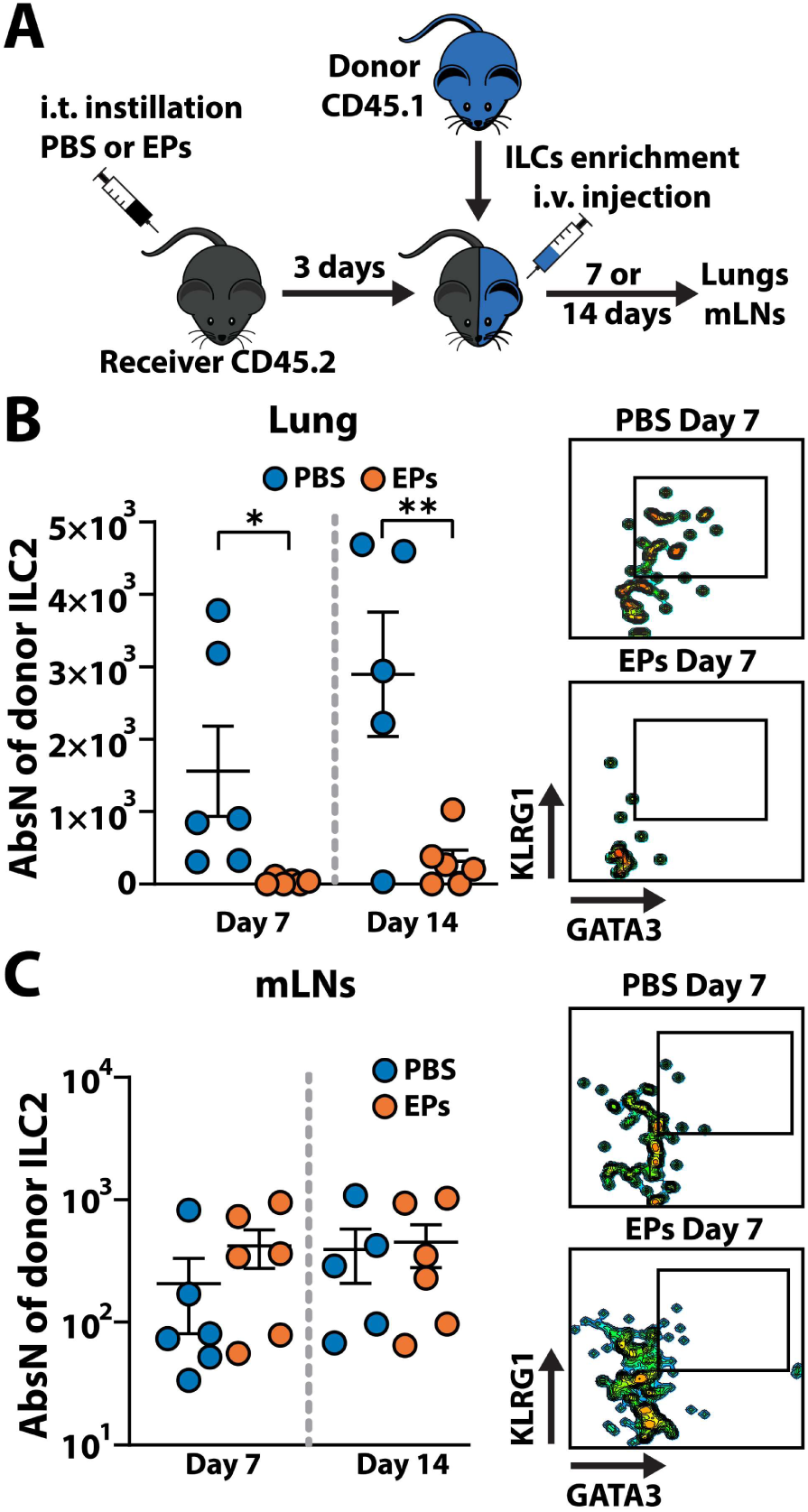
Analyses of the influence of EPs on ILC2 migration. **(A)** Experimental setup. Receiver CD45.2 mice were intratracheally instilled with 10 μg EPs in 50 μL PBS or with 50 μL PBS alone. 3 days after, ILCs from spleen and lymph nodes of CD45.1 donor mice were enriched by magnetic beads and adoptively transferred to the receiver mice. A ratio of 3 donors per receiver was used. 7 or 14 days after, mice were exsanguinated by cardiac perfusion, lungs and mediastinal lymph nodes were harvested and analyzed by flow cytometry. Absolute number of donor derived ILC2 7 (PBS n=6, EPs n=6) and 14 days (PBS n=5, EPs n=6) after transfer (left panel), and representative flow cytometry plots (right panel) in **(B)** lungs and **(C)** mediastinal lymph nodes. Data are presented as scatter plots with mean ± SEM. Differences between two groups were evaluated by using the Wilcoxon-Mann–Whitney test. P values <0.05 were considered significant.

To determine whether this effect extends to a stronger inflammatory context, we used a model of acute house dust mite (HDM)–induced lung inflammation, known to triggers robust ILC2 recruitment (*24*). Mice were immunized with HDM and challenged intratracheally with HDM alone or HDM plus EPs (Fig. 3A). EP-treated mice showed a pronounced reduction in lung ILC2 numbers, with no change in mLNs ILC2s and a trend toward increased circulating ILC2s (Fig. 3B–C). EPs did not alter the recruitment of GATA3⁺ CD4⁺ T cells or mast cells but significantly reduced eosinophil accumulation (Fig. 3D). Together, these findings establish that EPs prevent ILC2 trafficking to the lung across inflammatory settings.

**Figure 3:**
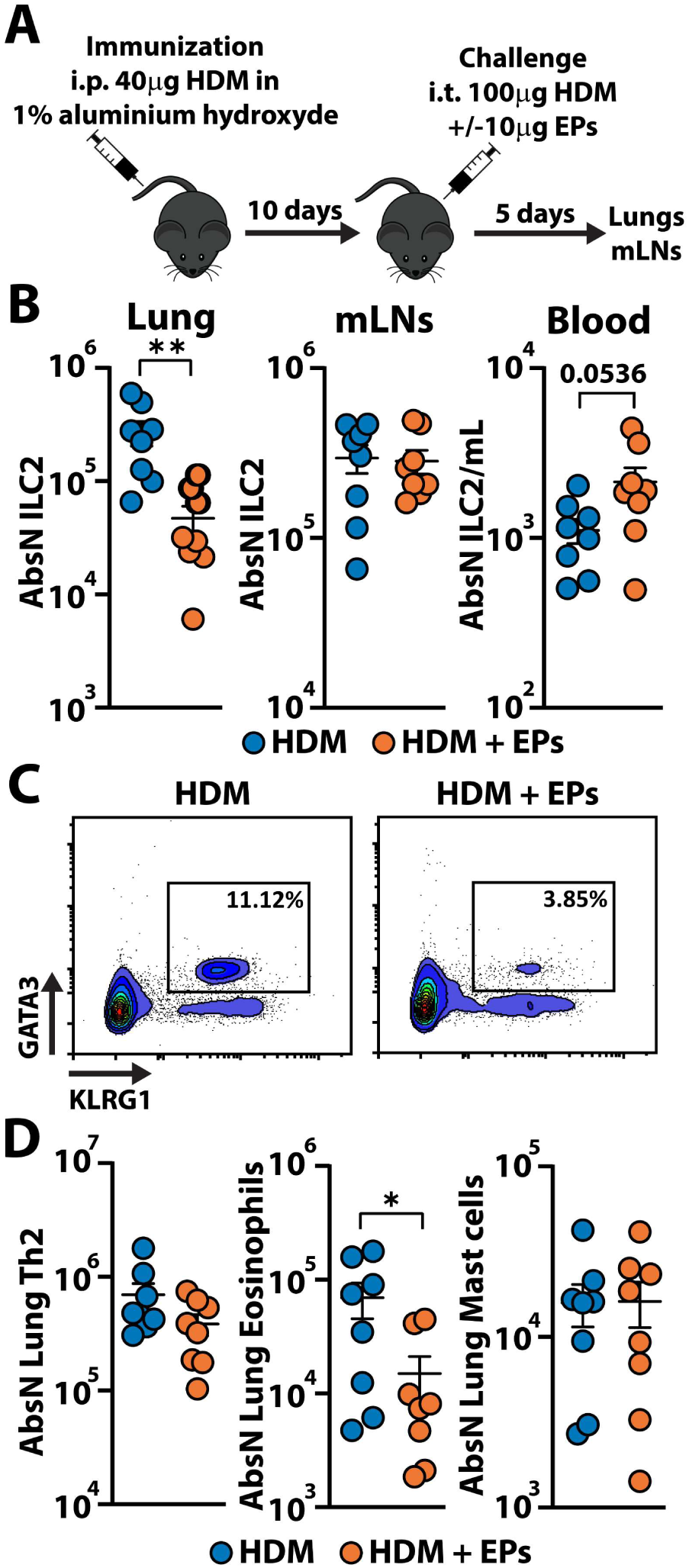
Impact of EPs on ILC2 recruitment in house-dust mite extract induced lung inflammation. **(A)** Experimental setup. Mice were immunized by an i.p. injection of 40 μg of HDM extract in 1% aluminum hydroxide. 10 days later, mice are challenged with an intra-tracheal instillation of 100 μg of HDM extract with or without 10 μg of EPs. 5 days later, lungs, mediastinal lymph nodes and blood were harvested and analyzed by flow cytometry. **(B)** Absolute number of ILC2 in the lungs, mediastinal lymph nodes and per mL of blood in mice challenged with HDM extract alone (n=8) or HDM extract + EPs (n=8). **(C)** Representative flow cytometry plots from lungs treated with HDM extract alone or HDM extract + EPs. **(D)** Absolute number of Th2 CD4 T cells, eosinophils and mast cells in the lungs of mice challenged with HDM extract alone (n=8) or HDM extract + EPs (n=8). Data are presented as scatter plots with mean ± SEM. Differences between two groups were evaluated by using the Wilcoxon-Mann–Whitney test. P values <0.05 were considered significant.

### EPs suppress CCL20 expression to prevent ILC2 recruitment

To investigate how EPs impair ILC2 recruitment to the lung, we first analyzed publicly available single-cell RNA-sequencing data comparing lungs from patients with chronic obstructive pulmonary disease (COPD) and healthy controls (*25*), a relevant human disease characterized by elevated levels of EPs (*16*, *26*). Gene ontology analysis revealed a coordinated down-regulation of pathways related to extracellular matrix organization and cell migration in COPD lungs (Fig. 4A), a pattern also observed within alveolar macrophages, monocyte/macrophage populations, and dendritic cells (Fig. S10) after clustering and identification (Fig. 4B). Among the most strongly down-regulated genes across these cell types was CCL20, both at the whole-lung level (Fig. 4C) and within individual myeloid subsets (Fig. 4D–E).

**Figure 4:**
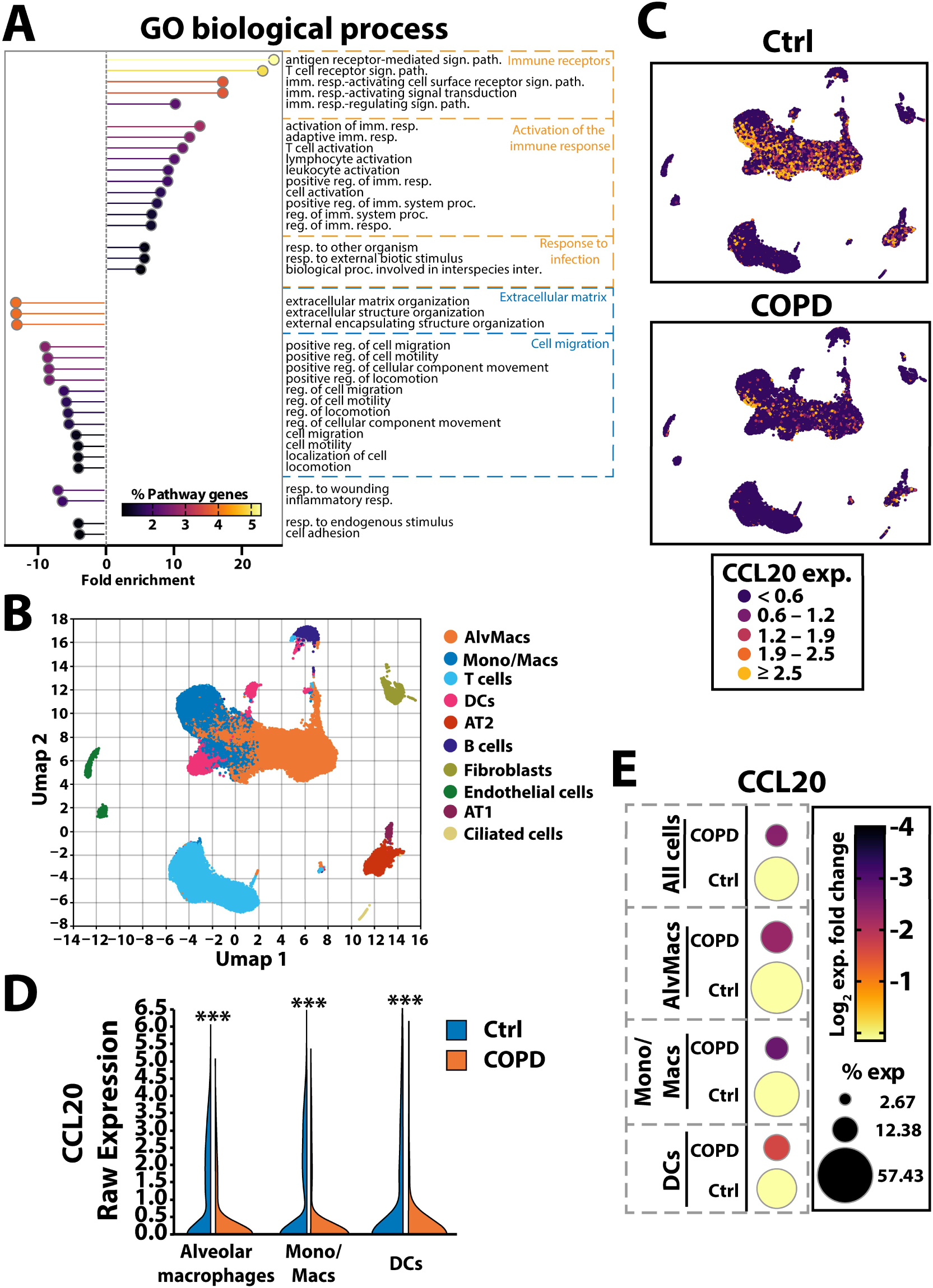
Analysis of single-cell RNA sequencing data from the lungs of chronic obstructive pulmonary disease or healthy lungs. Public scRNAseq data from (*25*) were analyzed. **(A)** Gene Ontology biological process up and downregulated in patients with COPD, from differential gene expression analysis. **(B)** UMAP dimensional reduction and cluster identification of the different cell populations. **(C)** Heatmap of CCL20 expression on UMAP from healthy lungs (left) or in lungs of patients with COPD (right). **(D)** Raw expression of CCL20 in specific cell clusters comparing healthy lungs (blue) and COPD patients’ lungs (orange). **(E)** Fold change in CCL20 expression comparing healthy lungs and COPD patients’ lungs in all cells or in myeloid cell clusters.

We next quantified CCL20 in the serum of patients with COPD at stable state and healthy donors (Table S1) and found that CCL20 levels correlated with the proportion of circulating ILC2s in both patients with COPD and healthy donors (Fig. 5A; Fig. S11). Lower serum CCL20 concentrations were associated with clinical features linked to higher EPs burden (Fig. 5; Fig. S12), including emphysema at diagnosis (Fig. 5B), higher hyperinflation with increased residual volume and total lung capacity (Fig. 5C–D), recent acute exacerbations (Fig. 5E), and a trend toward heavier smoking (Fig. 5F). Together, these data indicate that CCL20 expression is reduced in clinical settings characterized by elevated EPs and that CCL20 levels track with ILC2 abundance.

**Figure 5:**
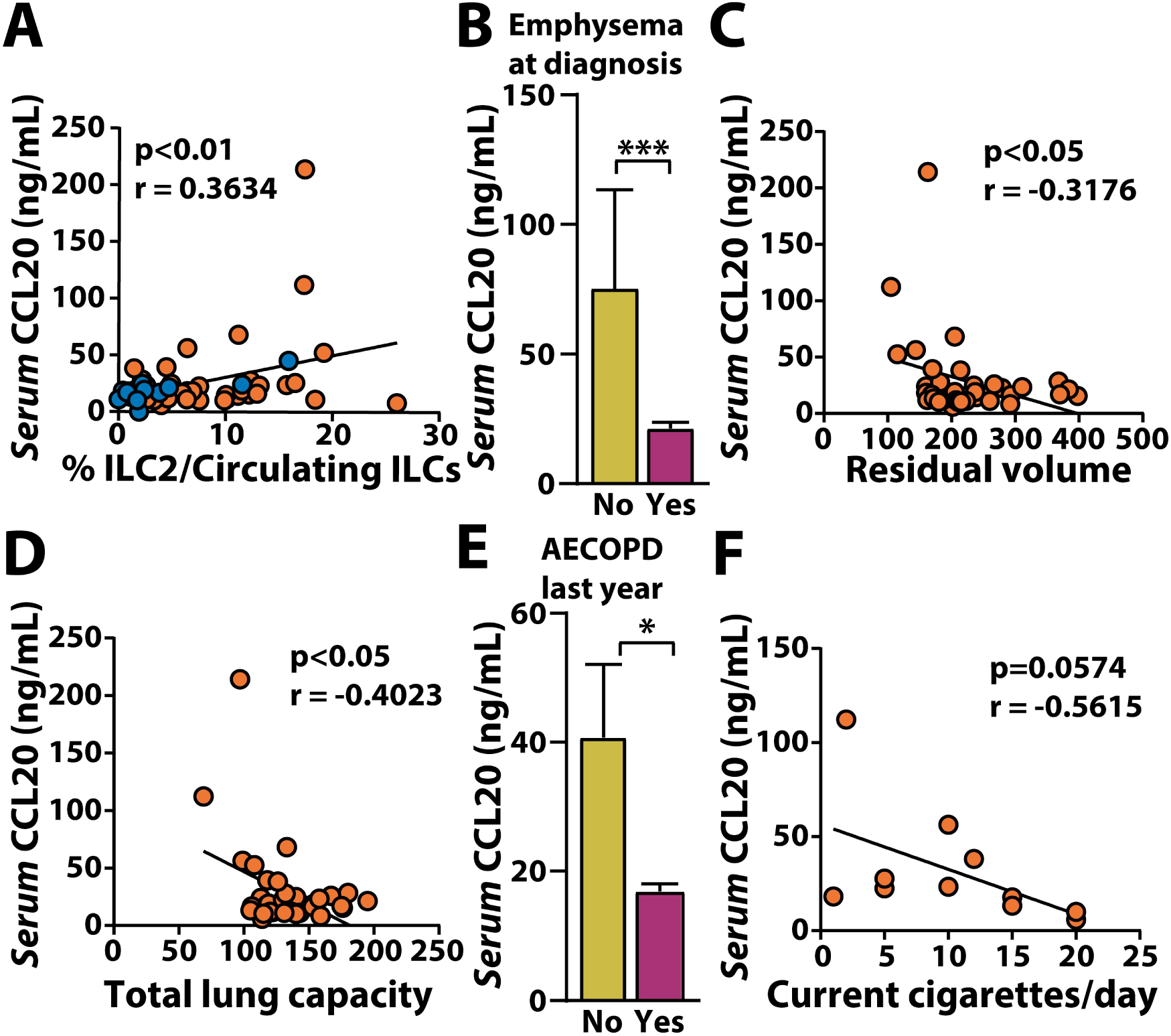
CCL20 serum levels in patients with COPD and correlation to clinical factors. Blood was drawn from patients with COPD and healthy donors, PBMCs were analyzed by flow cytometry and serum was collected for CCL20 quantification by cytometric bead assay. **(A)** Correlation between serum level of CCL20 and percentage of circulating ILC2. Blue dots indicate healthy donor and orange dots patients with COPD. **(B-F)** CCL20 serum level according to emphysema at diagnosis **(B)**, lung residual volume **(C)**, total lung capacity **(D)**, disease exacerbation during the last year **(E)** and current smoking **(F)** in patients with COPD. Data in bar graphs are presented as mean ± SEM. Differences between two groups were evaluated by using the Wilcoxon-Mann–Whitney test. For correlations Pearson r test was used. P values <0.05 were considered significant.

To directly test whether EPs suppress CCL20 during lung inflammation, we returned to the HDM-induced inflammation model. EPs treatment did not alter CCR6 expression—the receptor for CCL20—on lung ILC2s (Fig. 6A–B), indicating that impaired recruitment is not due to defective receptor availability. Instead, EPs markedly reduced the proportion of lung cells expressing CCL20 (Fig. 6C–D) and noticeably decreased CCL20 signal intensity at the single-cell level (Fig. 6E–H). These findings demonstrate that EPs suppress CCL20 production in the inflamed lung, thereby blocking ILC2 recruitment.

**Figure 6:**
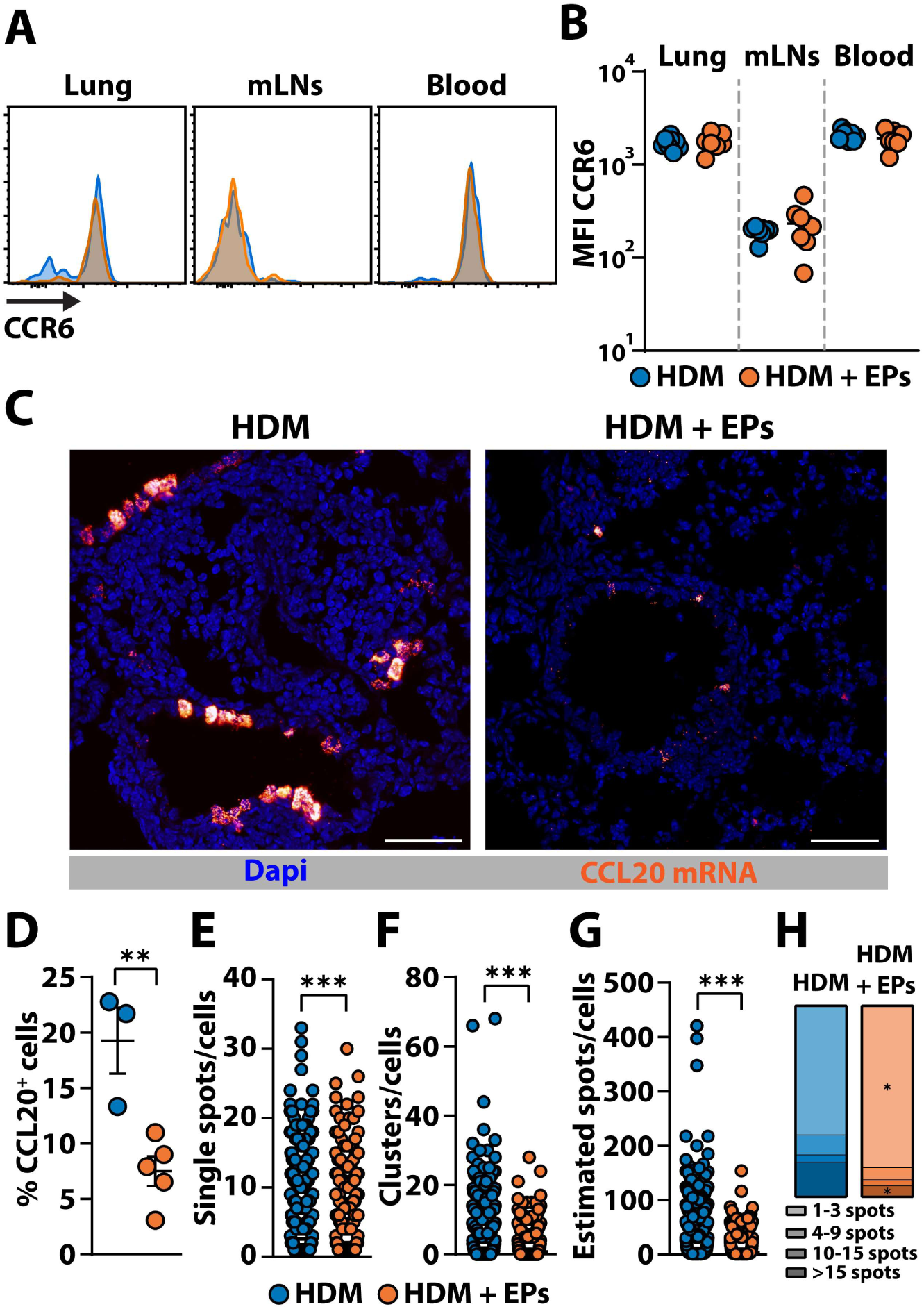
Analysis of the influence of EPs on CCL20 and its receptor CCR6 in house-dust mite extract induced lung inflammation. Representative flow cytometry histograms **(A)** and quantification **(B)** of CCR6 expression on ILC2 from lungs, mediastinal lymph nodes and blood of mice treated with HDM extract alone (blue) or HDM extract + EPs (orange). **(C)** Representative photomicrographs of fluorescence *in situ* hybridization of CCL20 (orange hot) in the lung of HDM (left) or HDM+EPs (right) treated mice. Tissue sections were counterstain with dapi (blue) to identify nuclei. Scale bar = 50μm. **(D)** Quantification of CCL20 mRNA expression. **(D)** Proportion of CCL20 expression in cells. **(E)** Number of single spots per expressing cells. **(F)** Number of clusters per expressing cells. **(G)** Estimated number of spots per expressing cells after cluster segmentation. **(H)** Scores of expression. Data in bar graphs are presented as mean ± SEM. Differences between two groups were evaluated by using the Wilcoxon-Mann–Whitney test. P values <0.05 were considered significant.

## DISCUSSION

Our study identifies elastin-derived peptides as previously unrecognized regulators of ILC2 recruitment during lung inflammation. Although EPs have long been known to accumulate in chronic lung disease and to modulate diverse immune populations (*20*, *22*, *23*), their impact on innate lymphoid cells had not been defined. We show here that EPs selectively prevent entry of circulating ILC2 in the lung by suppressing CCL20 expression upon inflammation, revealing a direct link between ECM degradation and the chemokine networks that orchestrate type 2 immunity.

ILC2 were long viewed as a largely tissue-resident and relatively homogeneous population (*27–29*). However, helminth infection revealed the existence of circulating IL-25–responsive inflammatory ILC2 (iILC2) that migrate to the lung (*6*). These cells arise from an intestinal pool (*7*, *30*) and transit through lymphatic vessels and mesenteric lymph nodes before entering the bloodstream and reaching the lung (*31*, *32*). While S1P-mediated tissue egress is essential for their mobilization (*6*, *7*), the mechanisms guiding their recruitment from the circulation into the lung remain poorly defined but are likely chemokine-driven. Here, we identify CCL20 as a key chemokine guiding ILC2 recruitment during acute lung inflammation. CCL20 is classically associated with CCR6⁺ lymphocytes and myeloid cells (*33*). Its expression is induced by HDM and cigarette smoke (*34*, *35*) and is elevated in the sputum of asthmatic and patients with COPD (*36*). CCL20 also promotes the recruitment of regulatory T cells (Tregs) during allergic inflammation and tobacco exposure (*37*, *38*), raising the possibility that EPs may influence Treg positioning. Whether this effect dampens inflammation or promotes pathogenic Th17 conversion (*39*) remains to be determined. EP-mediated suppression of CCL20 almost completely abrogated ILC2 recruitment without altering T cell infiltration, indicating that alternative inflammatory chemokines cannot compensate for the loss of CCL20. This was unexpected given prior reports implicating CCR9/CCL25 and CXCR6/CXCL16 in ILC2 migration (*40*, *41*), neither of which was differentially regulated in COPD lungs in our dataset. CCR4 has also been reported on IL-25–induced lung-recruited ILC2 (*42*), although its functional relevance remains unclear. The partial reduction of ILC2 recruitment in the HDM model suggests that CCR4-dependent cues may provide limited compensation in this context but are insufficient in EP-induced inflammation. Notably, CCL20 has recently been implicated in ILC2 recruitment in cystic fibrosis (*43*), a condition sometimes associated with emphysema (*44*, *45*). Moreover, gut- and mesenteric lymph node–derived ILC2, which give rise to iILC2, express CCR6 (*46*). Together, these observations suggest that reliance on CCL20 may be context-dependent and become dominant in microenvironments enriched in EPs.

Because elastin degradation is a hallmark of COPD, asthma, and other chronic lung diseases (*47*, *48*), EP-mediated suppression of CCL20 may represent a general mechanism by which tissue damage modulates type 2 immunity. This raises the possibility that ECM-derived signals contribute to disease heterogeneity by shaping the chemokine environment that governs ILC2 positioning. The relevance of this mechanism is supported by our analysis of patients with COPD, in whom elevated EP burden is a defining feature (*16*). We observed reduced CCL20 expression in COPD lungs and an inverse relationship between serum CCL20 levels and clinical indicators of elastin degradation, including emphysema severity and hyperinflation. These findings suggest that chronic EP exposure may contribute to impaired ILC2 recruitment in COPD and other chronic lung conditions. Further investigation is warranted to determine how this mechanism influences the balance between protective and pathological type 2 responses and whether it can be therapeutically targeted. Indeed, ILC2 contribute to epithelial repair (*49*) and protect against viral (*50*) and bacterial infections (*51*), but also promote fungal infections (*52*) and eosinophil-driven allergic inflammation (*53*). Fine-tuning their recruitment is therefore critical to disease outcome.

Asthma and allergic inflammation are dominated by pathogenic type 2 responses (*54*), making it unlikely that targeting EPs would be beneficial in this context. COPD, by contrast, is a mixed inflammatory disease (*55*) and a prototypical condition of high EP burden (*16*). ILC2 have been proposed to contribute to type 2 immunity in COPD due to their increased numbers and cytokine production (*21*, *56*, *57*), although whether they are protective or pathogenic remains unresolved. Elevated circulating ILC2 in acute exacerbations (*58*, *59*) may reflect impaired migration into the lung, consistent with our findings. ILC2 may also contribute to pathology in eosinophilic COPD, where inflammation and IL-33 levels are elevated (*60*). Our observation that decreased lung ILC2 correlates with reduced eosinophils supports this possibility. Cigarette smoking, the major risk factor for COPD (*61*), induces elastin breakdown (*62*) and IL-33 expression (*63*), yet paradoxically reduces IL-33 levels in patients with COPD who continue to smoke (*64*). Moreover, IL-33 blockade reduces exacerbations in former smokers but not in current smokers (*65*). In our study, serum CCL20 tended to decrease with higher cigarette consumption, suggesting that active smoking and elevated EP burden may impair type 2 immunity. This may reflect smoke-induced dysregulation of IL-33 signaling, which silences ILC2 repair functions and promotes exacerbations (*66*). Thus, targeting EPs during acute exacerbated COPD (AECOPD) to restore ILC2 migration to the lung may represent a promising therapeutic strategy.

The limitations of our study are that although we identify CCL20 suppression as a key mechanism by which EPs impair ILC2 recruitment, the upstream signaling pathways through which EPs regulate CCL20 expression remain to be defined. In addition, while our human data support a link between EP burden and reduced CCL20 levels, causality cannot be established in this setting.

Overall, our work establishes EPs as regulators of CCL20-dependent ILC2 recruitment and reveals a pathway linking ECM breakdown to altered type 2 immunity. These findings broaden the understanding of ECM–immune crosstalk and suggest new strategies for modulating immune responses in chronic lung disease.

## MATERIALS AND METHODS

### Animal procedures

#### Mice

6 weeks female C57BL/6J mice or B6.Cg-Ptprc^a^-Pepc^b^/Rj (carrying the CD45.1 allele) were purchased from Charles River or bred in-house. Mice were housed in the same controlled environment (12h-light-dark cycle, 22 ± 2°C, 30–70% humidity), with standard rodent chow and water ad libitum. All procedures and protocols conducted on mice were consented by the ethics committee 56 of Reims Champagne Ardenne University and by the French Ministry of Education and Research and they comply with institutional guidelines and regulations relating to animal research. In all experiments, euthanasia was performed by an overdose of sodium pentobarbital (120 mg/kg) under anesthesia and followed by an intracardiac perfusion with PBS 5mM EDTA. Lung, mediastinal lymph nodes (mLNs) and peripheral lymph nodes (pLNs = inguinal, axillar and brachial lymph nodes) were collected. Specifically for experiments deciphering ILC2 migration, a blood tissue partitioning method (2 minutes before euthanasia, mice were i.v. injected with 2 mg of anti-CD45 APC to label blood cells) was used instead of perfusion.

#### Intratracheal instillations

Mice were anesthetized with a mixture of ketamine (80 mg/kg) and xylazine (10 mg/kg) administrated intraperitoneally. Intratracheal instillation was performed using an open otoscope adapted for use as a laryngoscope. To ensure efficient delivery, mice were intubated with a 22G catheter into the oropharyngeal cavity. The volume for all procedures requiring intratracheal instillation is 50 μl.

#### Elastin peptides (EPs) induced lung inflammation model

For EPs induced lung inflammation, mice were intratracheally instilled with 10 mg of EPs or PBS vehicle. Mice were euthanized on day 3, 7, 14 or 21 after EPs instillation.

#### Adoptive transfer

Total innate lymphoid cells were enriched from spleen and lymph nodes of CD45.1 mice by negative selection using a modified version of the Dynabeads Untouched Mouse CD8 T Cells kit with anti-CD8 antibodies added to the enrichment cocktail. For experiments investigating the proliferation of ILCs, enriched cells were stained with CFSE before adoptive transfer. Briefly, cells were washed twice and stained with CFSE for 8 minutes at 37°C and 5% CO2. Reaction was stopped with cold FBS for 2 minutes on ice and cells were washed twice before resuspension in PBS. Enriched ILCs were transferred to recipient CD45.2 mice by intra-venous injection in a volume of 100 μl of PBS.

#### House dust mite (HDM) allergy model

Mice were sensitized to HDM by an intraperitoneal injection of 40 μg of extract (*D. pteronyssinus* mite bodies extract, Citeq) in PBS. Ten days after the first exposure to HDM extract, mice were challenged by an intratracheal instillation of 100 μg of HDM extract with or without 10 μg of EP. Five days after the challenge, mice were euthanized and lung, mLNs and blood were collected.

### Cell suspensions

#### Mouse cells

For the lungs, perfused lung mice were cut into small pieces and digested with a mixture of collagenase D (1 mg/ml) and DNase (0.1 mg/ml) in PBS FBS 10% for 20 minutes at 37°C under agitation. Lung homogenates were filtered through a 100 μm cell strainer, washed with RPMI (10% FBS and antibiotics) and centrifugated (450g, 5 minutes, 4°C). Leukocyte fraction was enriched by centrifugation in a 40/80% Percoll gradient (300g, 23 minutes, 20°C, acceleration: 5, brake: 5). Lymph nodes suspensions were obtained by mechanical disruptions of the lymph nodes and filtering through a 40 μm cell strainer. For all suspensions, cells were washed with PBS FBS 10% before being resuspended in PBS 5% FBS 0.1% NaN_3_ for antibody staining.

#### Human peripheral blood mononuclear cells (PBMC)

Whole blood from patients with COPD was collected in heparin/lithium tubes. The blood was transferred in a 50 mL tube, and the same volume of sterile PBS was added. 16 ml of Pancoll was gently added below diluted blood. After centrifugation (800g, 30 minutes, 20°C, without brake), the PBMC ring was recovered and washed with PBS (qsp 40 ml). Cells were resuspended in PBS 5% FBS 0.1% NaN_3_ for further analysis.

### Flow cytometry

Cell suspensions were obtained as previously detailed. Analysis of the surface and intracellular markers expression was performed using antibodies listed in Table 1. Surface staining was performed on 5.10^6^ cells in 20 μL of antibodies in PBS 5% FBS 0.1% NaN_3_ for 30 minutes under agitation at 4°C in the dark. Cells were then fixed and permeabilized with eBioscience transcription factor staining buffer set according to the manufacturer’s protocol. For intranuclear markers, incubation was performed overnight at 4°C. Cells were washed and resuspended in PBS and acquisition was performed using a BD LSRFortessa cell analyzer (BD Biosciences). Results were analyzed with Flowlogic v7.3 (Inivai Technologies) or FlowJo v10 softwares.

### Antibodies used in the study

**Table 1:** Antibodies used in the study.

| Antibody | Clone | Fluorochrome | Reference |
| --- | --- | --- | --- |
| Anti-mouse antibodies |  |  |  |
| CD19 | 6D5 | PE-Cy7 | Biolegend - 115520 |
| CD11b | M1/70 | PE-Cy7 | Biolegend - 101216 |
| CD11c | N418 | PE-Cy7 | Biolegend - 117318 |
| TcRgd | GL3 | PE-Cy7 | Biolegend - 118124 |
| TER119 | TER-119 | PE-Cy7 | Biolegend - 116222 |
| TcRb | H57-597 | PE-Cy7 | Biolegend - 109222 |
| TcRb | H57-597 | PerCP-Cy5.5 | BD Biosciences - 560657 |
| CD45 | 30F11 | BV711 | Biolegend - 103147 |
| CD45 | 30-F11 | APC | Biolegend - 103112 |
| CD45.1 | A20 | BV711 | Biolegend - 110739 |
| CD45.2 | REA1223 | PerCP-Vio770 | Miltenyi - 130-124-087 |
| CD45.2 | 104 | APC | Biolegend - 109814 |
| KLRG1 | 2F1/KLRG1 | PE | Biolegend - 138408 |
| NKp46 | 29A1.4 | A647 | Biolegend - 137628 |
| NKp46 | 29A1.4 | BV510 | Biolegend - 137623 |
| CD127 | A7R34 | BV785 | Biolegend - 135037 |
| Zombie NIR |  | APC/Cy7 | Biolegend - 423105 |
| CD4 | GK1.5 | BV510 | Biolegend - 100449 |
| CD8 | 53-6.7 | APC-vio770 | Miltenyi - 130-123-280 |
| RORyt | Q31-378 | BV650 | BD Biosciences - 564722 |
| T-bet | REA102 | PE-vio615 | Miltenyi - 130-121-998 |
| GATA-3 | 16E10A23 | BV421 | Biolegend – 653814 |
| Anti-human antibodies |  |  |  |
| Lineage | UCHT1; HCD14; 3G8;<br>HIB19; 2H7; HCD56 | Pacific Blue | Biolegend - 348805 |
| CD7 | M-T701 | PE-CF594 | BD Biosciences – 562541 |
| CD127 | HIL-7R-M21 | PE-Cy7 | BD Biosciences – 560822 |
| CD294/CRTH2 | BM16 | A647 | BD Biosciences – 558042 |
| CD117 | 104D2 | BV605 | BD Biosciences – 562687 |
| Tbet | AD2 | PE | BD Biosciences - 561258 |
| GATA3 | L50-823 | BB700 | BD Biosciences - 566642 |
| RORyt | REA278 | Vio-B515 | Miltenyi - 130-123-840 |

### Quantification of inflammatory cells by microscopy

Lungs from PBS- or EP-treated mice were fixed, paraffin-embedded, and sectioned at 3 µm. After antigen retrieval, sections were stained with anti-Gr1 followed by an A488-conjugated secondary antibody and counterstained with DAPI. Slides were mounted with antifade medium, imaged on a Zeiss LSM880 confocal microscope (20×), and analyzed using QuPath.

### Single cell RNA-seq processing

Publicly available scRNA-seq datasets from COPD and healthy lungs were analyzed (GEO accession number GSE136831). Data were processed in Trailmaker (Parse Biosciences) using standard quality-control, doublet filtering, normalization, Harmony integration, dimensionality reduction, UMAP embedding, and Louvain clustering. Differentially expressed genes (adjusted p < 0.05, average expression ≥ 2, log2FC ≥ 1.5) were identified in Trailmaker, and Gene Ontology analysis was performed using ShinyGO (https://bioinformatics.sdstate.edu/go/).

### CCL20 quantification in serum

Patients were recruited from a single-center study (NCT02924818) conducted in the service of Pulmonary Medicine at the University Hospital of Reims, under the approval of an ethics committee in human biomedical research (CPP Dijon EST I, No. 2016-A00242-49). All subjects gave their informed and written consent prior to inclusion in the study. Patients with COPD were enrolled in the study based on clinical assessments with a forced expiratory volume in 1-s (FEV_1_)/forced vital capacity (FVC) < 0.7 after bronchodilation. At inclusion, all patients were stable for at least 4 wk. Patients with asthma, allergic disease, tuberculosis, neoplasia, or other chronic respiratory diseases were excluded. Control subjects were recruited from the French Blood Institution and did not present any acute or chronic respiratory disease. Serums from patients with COPD or healthy donors were collected from the blood in dry tubes at the same time as the blood used for PBMC, retrieved after centrifugation for 10 minutes at 1200g at 20°C and stored at −20°C until processing. CCL20 levels were measured using a specific LEGENDplex^TM^ (Biolegend) kit, according to manufacturer’s instructions. Acquisition was performed on BD LSRFortessa cell analyzer (BD Biosciences) and analyzed by the dedicated Qognit software (Biolegend).

### Fluorescence *in situ* hybridization

Lungs from HDM- or HDM+EP-treated mice were fixed in PLP, cryoprotected in sucrose, embedded in OCT, and sectioned at 10 µm. CCL20 mRNA was detected using the RNAscope Multiplex Fluorescent V2 assay (ACD) with the CCL20 probe, following the manufacturer’s instructions. Signal amplification was performed with TSA Vivid Fluorophore 570, and nuclei were counterstained with DAPI. Images were acquired on a Zeiss LSM880 confocal microscope and analyzed using QuPath.

### Statistical analysis

Data are presented as mean ± SEM. Data analysis was performed using Prism 10 (GraphPad). Comparisons were performed either with Wilcoxon-Mann–Whitney test or Pearson r test as referenced in figure legends. P values inferior to 0.05 were considered significant with *<0.05, **<0.01 and ***<0.001.

## Supplementary Materials

Figs. S1 to S12.

## Acknowledgments

The authors would like to acknowledge Dr Sandra Audonnet from the URCACyt core facility, Dr. Christine Terryn from the PICT core facility and Mr. Anthony Pigeon Anthony Chauvin from the URCAnim core facility for technical assistance.

## Author contributions

Conceptualization: RLN, AP

Methodology: SL, CF, LP, AP

Investigation: SL, CF, LP, AP

Formal analysis: SL, AP

Resources: JMPC, GD, SLJ

Data curation: SL, JMPC, GD, AP

Visualization: SL, CF, AP

Funding acquisition: SP, SLJ, AP

Project administration: RLN, AP

Supervision: SP, RLN, AP

Writing – original draft: SL, CF, LP, AP

Writing – review & editing: SL, CF, LP, JMPC, GD, SLJ, SP, RLN, AP

## Data and materials availability

All data are available in the main text or the supplementary materials.

