## Supplementary information for "Elastin-derived peptides suppress CCL20 expression and block ILC2 recruitment during lung inflammation"

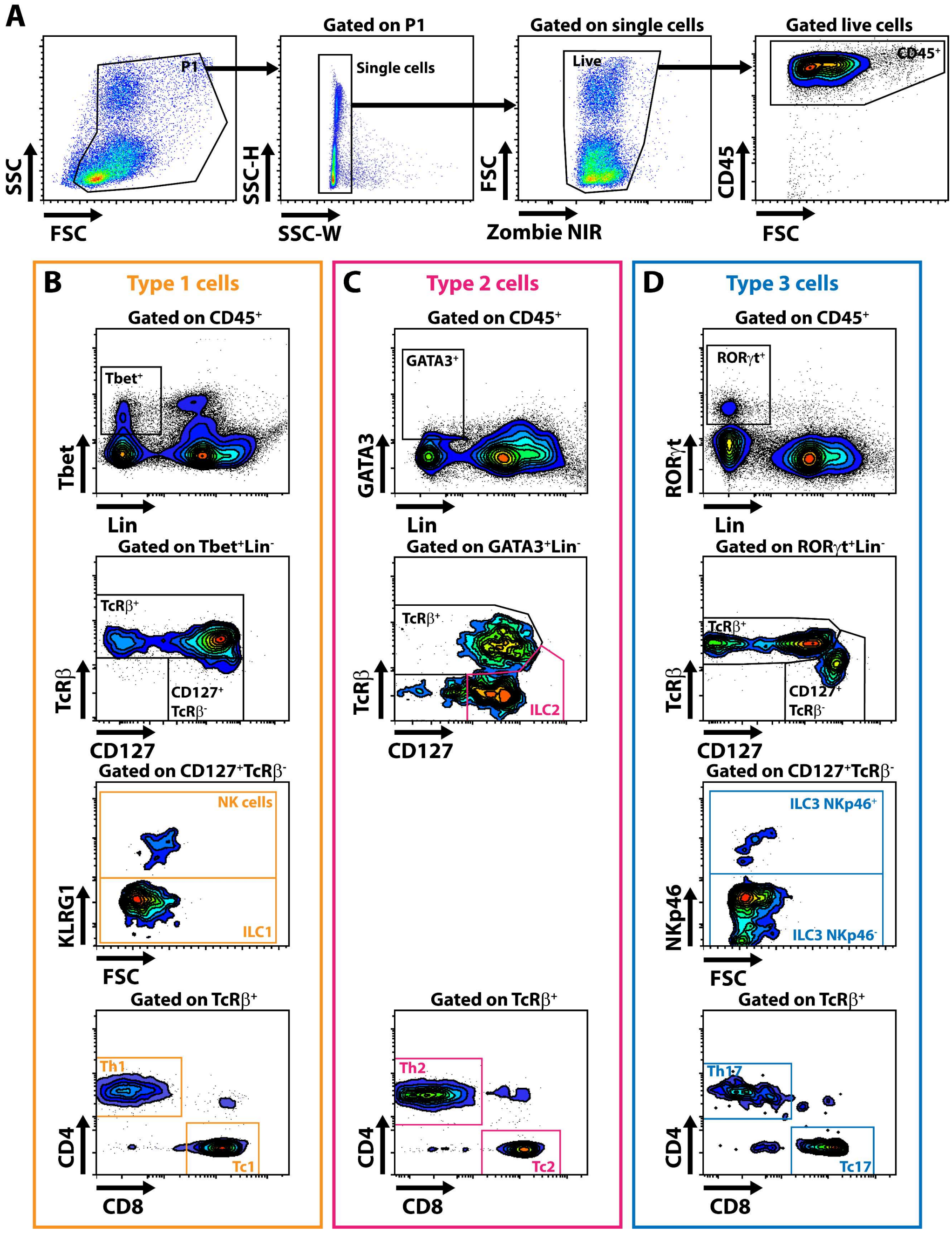


**Figure S1: Flow cytometry gating strategy used for the identification of ILCs and T cells subpopulations in the lungs of mice. (A)** Common gating to identify $CD45^+$ immune cells. **(B)** Gating of type 1 ILCs (ILC1), NK cells, Th1 and Tc1 cells. **(C)** Gating of type 2 ILCs (ILC2), Th2 and Tc2 cells. **(D)** Gating of NKp46 positive and negative type 3 ILCs (ILC3), Th17 and Tc17 cells.

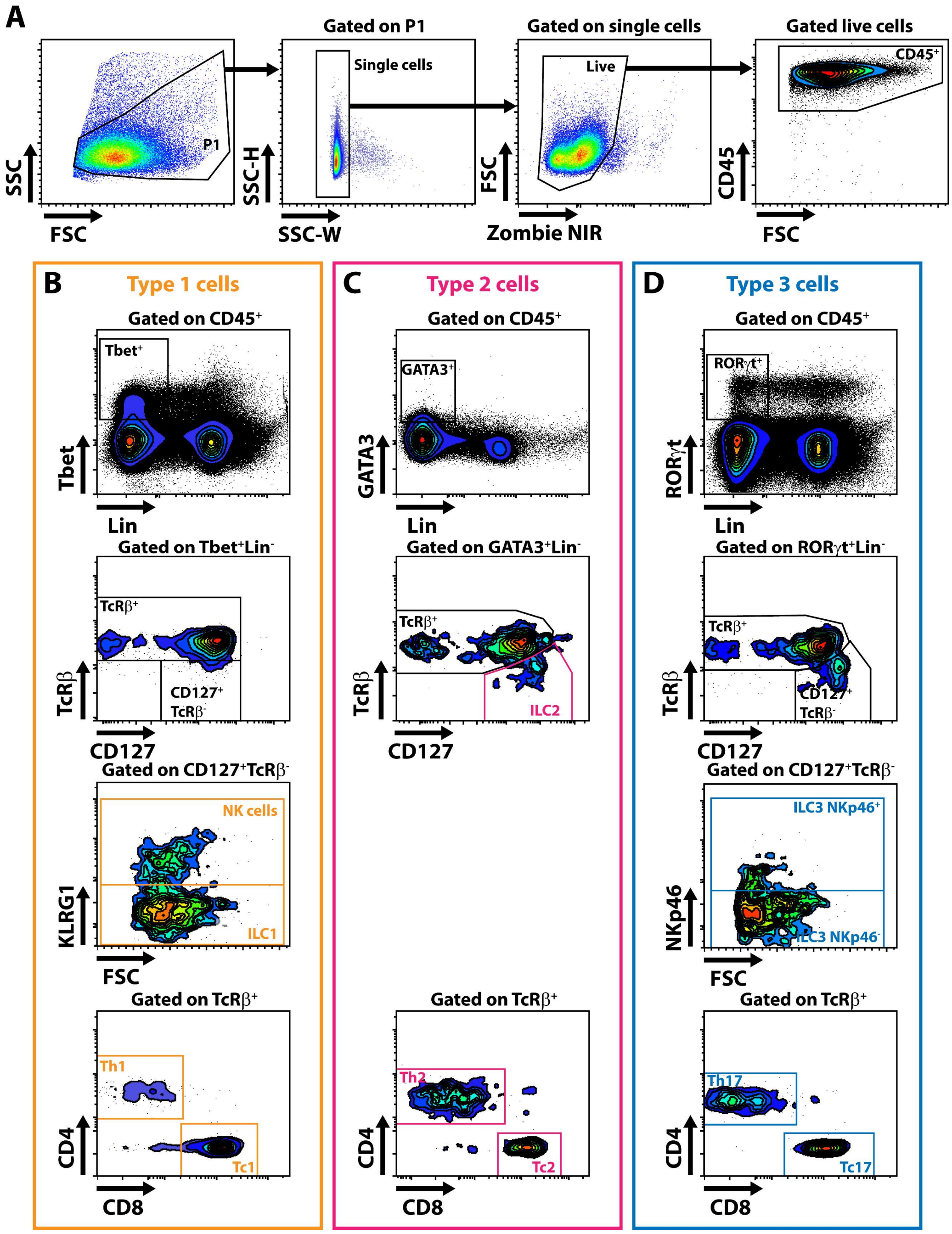


**Figure S2: Flow cytometry gating strategy used for the identification of ILCs and T cells subpopulations in lymph nodes. (A)** Common gating to identify CD45$^{+}$ immune cells. **(B)** Gating of ILC1, NK cells, Th1 and Tc1 cells. **(C)** Gating of ILC2, Th2 and Tc2 cells. **(D)** Gating of NKp46 positive and negative ILC3, Th17 and Tc17 cells.

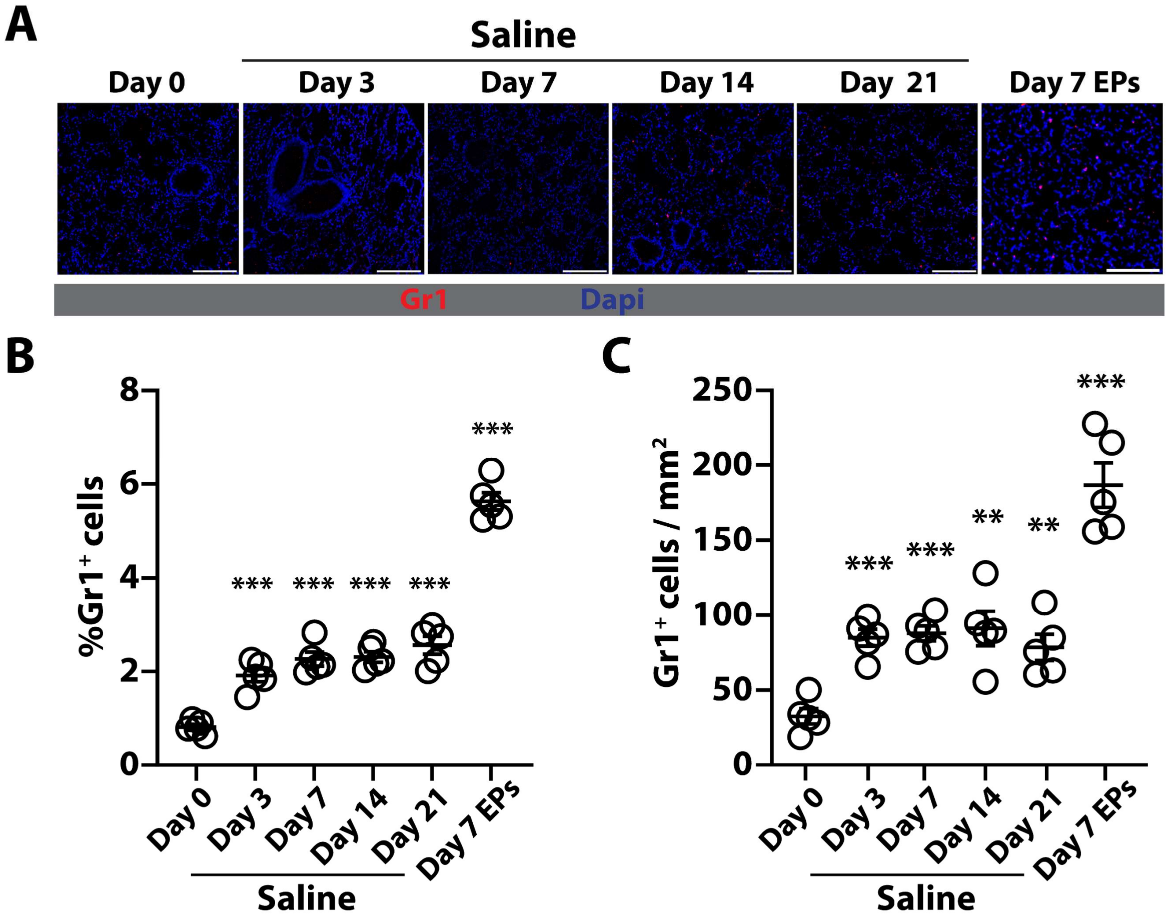


**Figure S3: Infiltration of inflammatory cells in the lung.** Lung sections from mice at basal state or after PBS (days 3, 7, 14 and 21) or EPs (Day 7) instillation were stained for Gr1 to identify inflammatory cells infiltration. **(A)** Representative micrographs of each condition. Quantification of **(B)** the percentage of $Gr1^{+}$ cells or **(C)** the number of $Gr1^{+}$ cells per mm2. N = 5 mice per condition. Data are presented as scatter plots with mean±SEM. Differences between day 0 and each other are indicated and were evaluated using the Wilcoxon-Mann–Whitney test. P values <0.05 were considered significant.

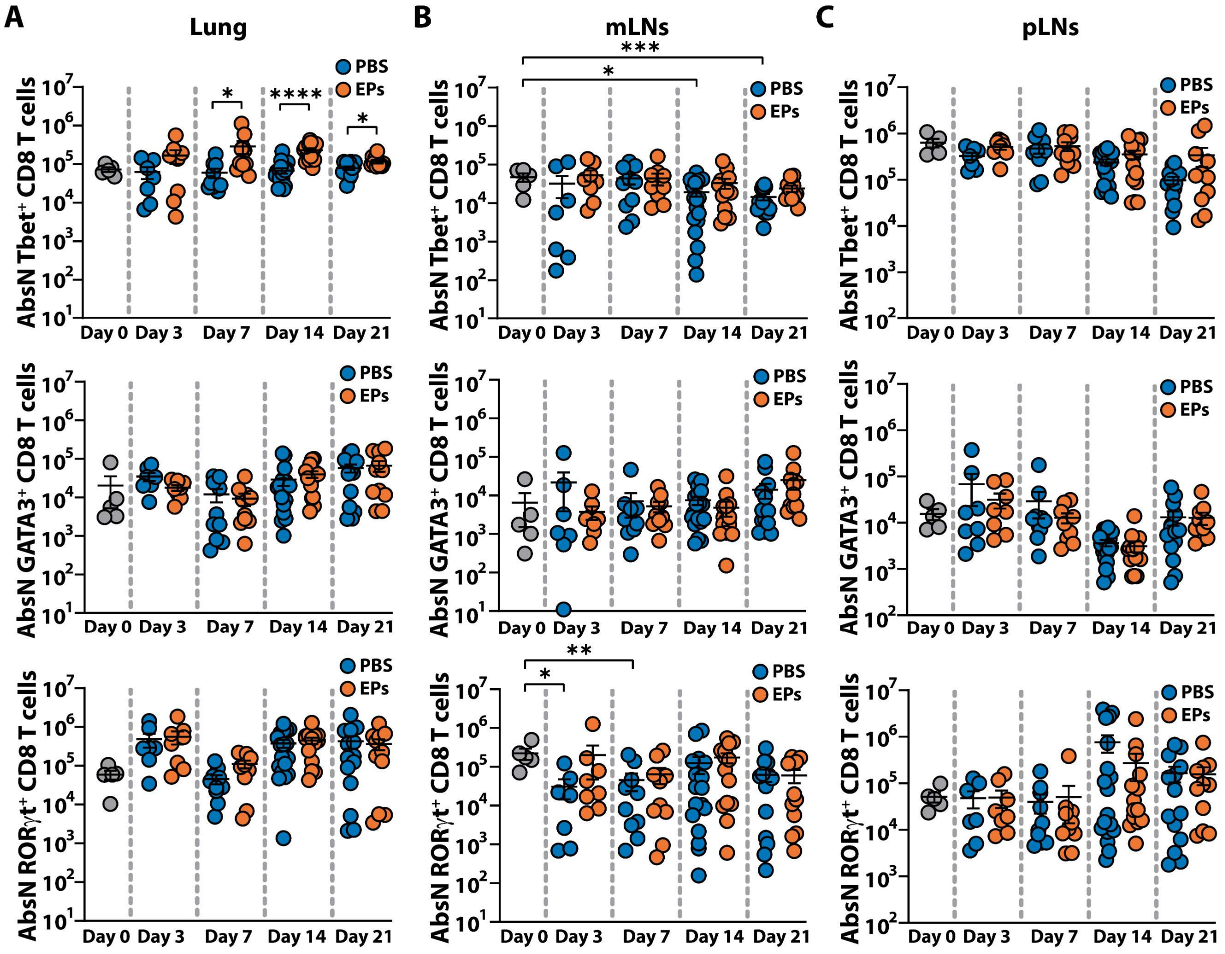


**Figure S4: Lungs, mediastinal and peripheral lymph nodes CD8 T cells numbers after EPs treatment.** Mice were intratracheally instilled with EPs or PBS and analyzed 0 (n=10), 3 (PBS n=10, EPs n=10), 7 (PBS n=16, EPs n=17), 14 (PBS n=15, EPs n=17) or 21 (PBS n=19, EPs n=18) days after. **(A)** Absolute number (AbsN) of lung Tbet$^+$ CD8 T cells (top panel), GATA3$^+$ CD8 T cells (middle panel) and RORγt$^+$ CD8 T cells (bottom panel). **(B)** AbsN of mediastinal lymph nodes Tbet$^+$ CD8 T cells (top panel), GATA3$^+$ CD8 T cells (middle panel) and RORγt$^+$ CD8 T cells (bottom panel). **(C)** AbsN of peripheral lymph nodes Tbet$^+$ CD8 T cells (top panel), GATA3$^+$ CD8 T cells (middle panel) and RORγt$^+$ CD8 T cells (bottom panel). Data are presented as scatter plots with mean±SEM. Differences between two groups were evaluated by using the Wilcoxon-Mann–Whitney test. P values <0.05 were considered significant.

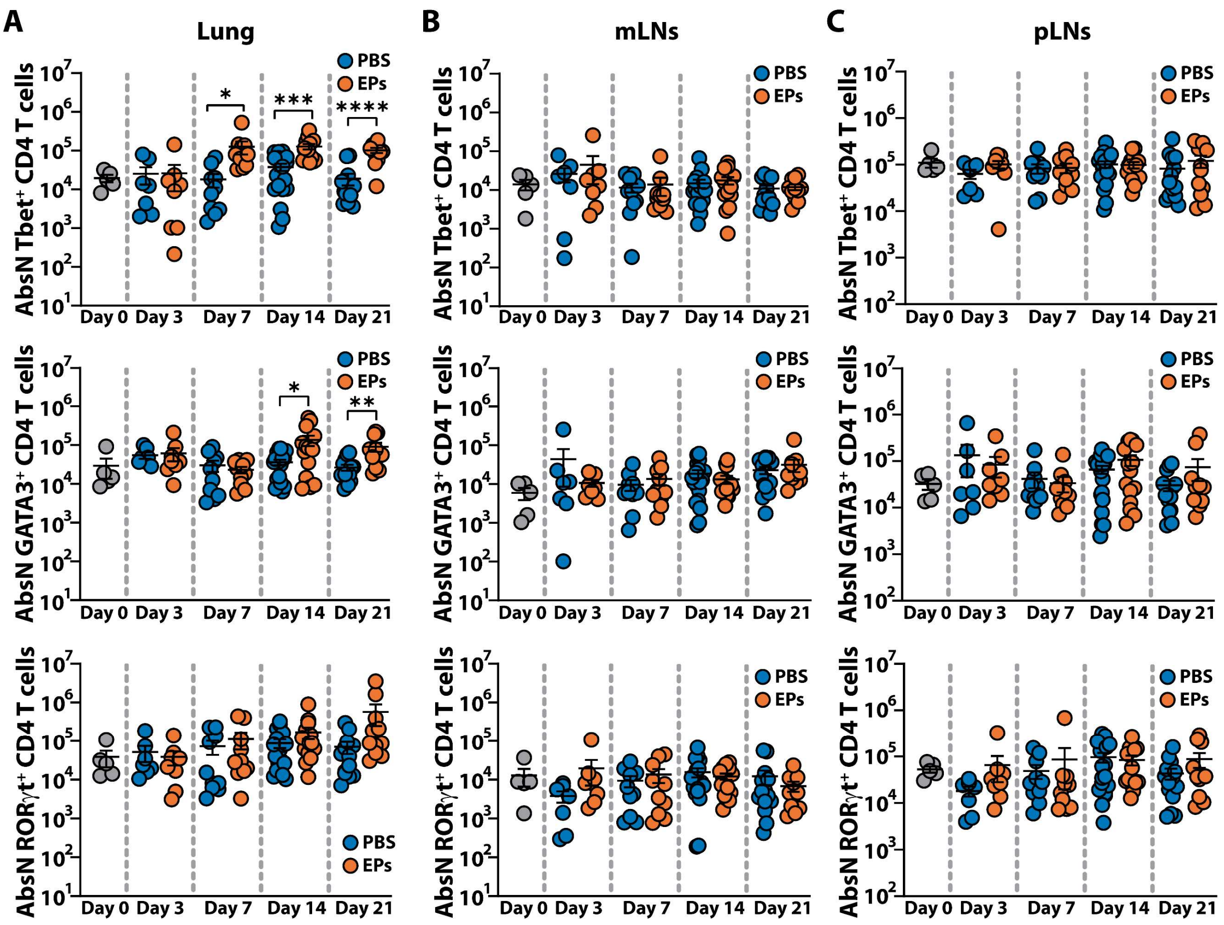


**Figure S5: Lungs, mediastinal and peripheral lymph nodes CD4 T cells numbers after EPs treatment.** Mice were intratracheally instilled with EPs or PBS and analyzed 0 (n=10), 3 (PBS n=10, EPs n=10), 7 (PBS n=16, EPs n=17), 14 (PBS n=15, EPs n=17) or 21 (PBS n=19, EPs n=18) days after. **(A)** AbsN of lung Tbet$^{+}$ CD4 T cells (top panel), GATA3$^{+}$ CD4 T cells (middle panel) and RORγt$^{+}$ CD4 T cells (bottom panel). **(B)** AbsN of mediastinal lymph nodes Tbet$^{+}$ CD4 T cells (top panel), GATA3$^{+}$ CD4 T cells (middle panel) and RORγt$^{+}$ CD4 T cells (bottom panel). **(C)** AbsN of peripheral lymph nodes Tbet$^{+}$ CD4 T cells (top panel), GATA3$^{+}$ CD4 T cells (middle panel) and RORγt$^{+}$ CD4 T cells (bottom panel). Data are presented as scatter plots with mean±SEM. Differences between two groups were evaluated by using the Wilcoxon-Mann–Whitney test. P values <0.05 were considered significant.

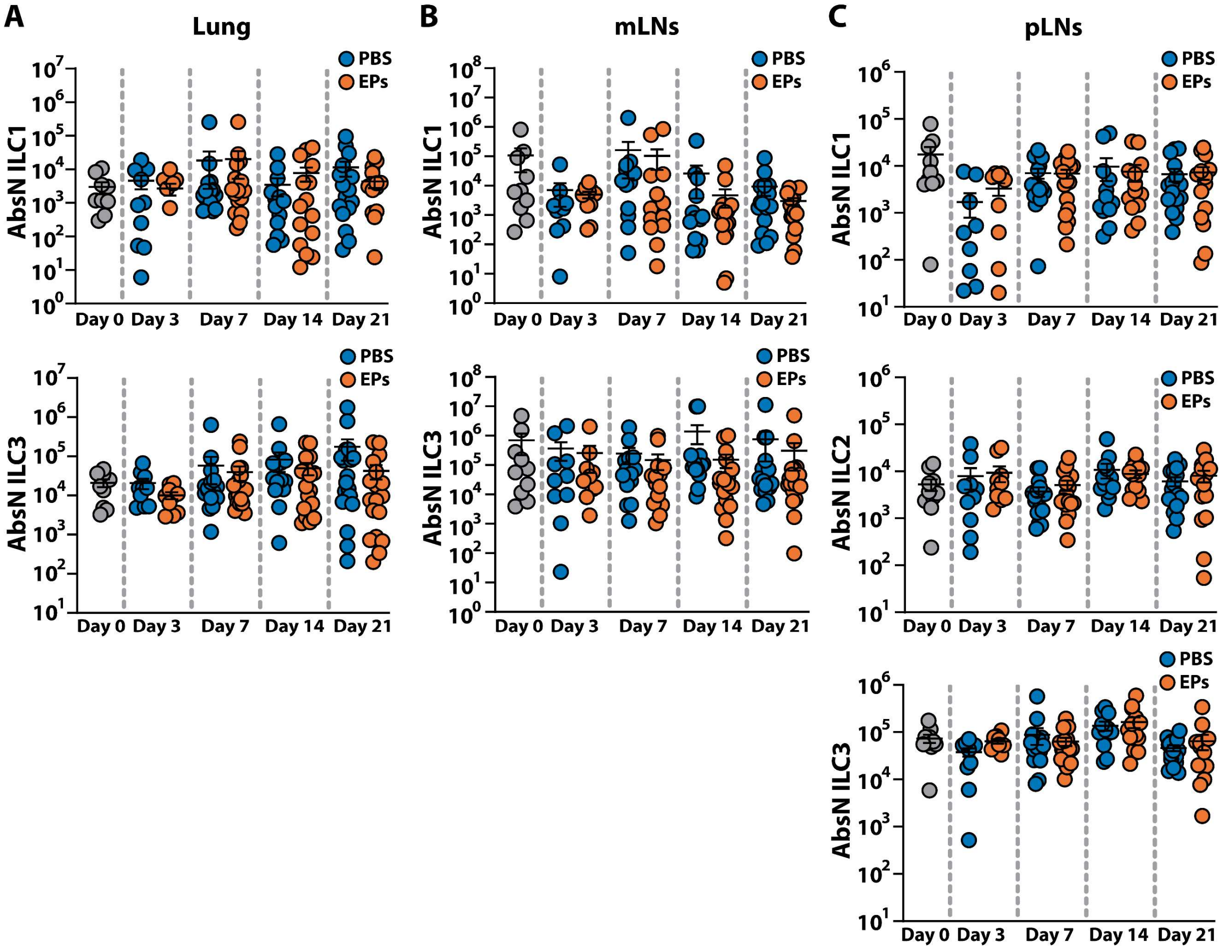


**Figure S6: Lungs, mediastinal and peripheral lymph nodes ILCs numbers after EPs treatment.** Mice were intratracheally instilled with EPs or PBS and analyzed 0 (n=10), 3 (PBS n=10, EPs n=10), 7 (PBS n=16, EPs n=17), 14 (PBS n=15, EPs n=17) or 21 (PBS n=19, EPs n=18) days after. **(A)** AbsN of lung ILC1 (top panel) and ILC3 (bottom panel). **(B)** AbsN of mediastinal lymph nodes ILC1 (top panel) and ILC3 (bottom panel). **(C)** AbsN of peripheral lymph nodes ILC1 (top panel), ILC2 (middle panel) and ILC3 (bottom panel). Data are presented as scatter plots with mean±SEM. Differences between two groups were evaluated by using the Wilcoxon-Mann–Whitney test. P values <0.05 were considered significant.

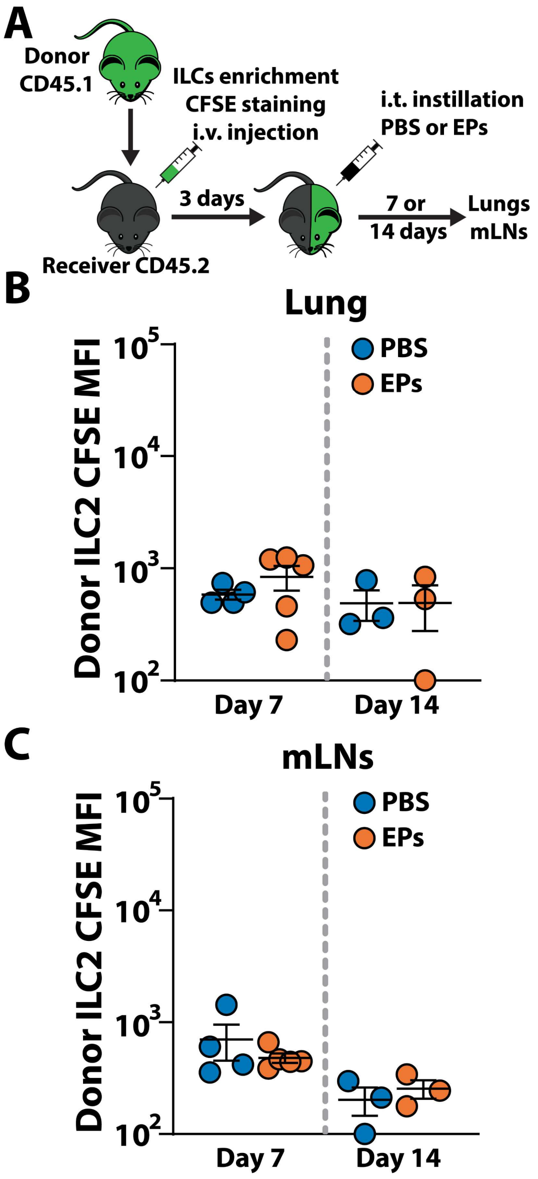


**Figure S7: Analyses of ILC2 proliferation. (A)** Experimental setup. ILCs from spleen and lymph nodes of CD45.1 donor mice were enriched by magnetic beads, CFSE stained and adoptively transferred to the CD45.2 receiver mice. 3 days after, receiver mice were intratracheally instillated with 10 µg EPs in 50 µL PBS or with 50 µL PBS alone. A ratio of 3 donors per receiver was used. 7 or 14 days after, mice were exsanguinated by cardiac perfusion, lungs and mediastinal lymph nodes were harvested and analyzed by flow cytometry. Mean fluorescence intensity of CFSE in ILC2 7 (PBS n=4, EPs n=5) and 14 days (PBS n=3, EPs n=3) after instillation in **(B)** lungs and **(C)** mediastinal lymph nodes. Data are presented as scatter plots with mean±SEM. Differences between two groups were evaluated by using the Wilcoxon-Mann–Whitney test.

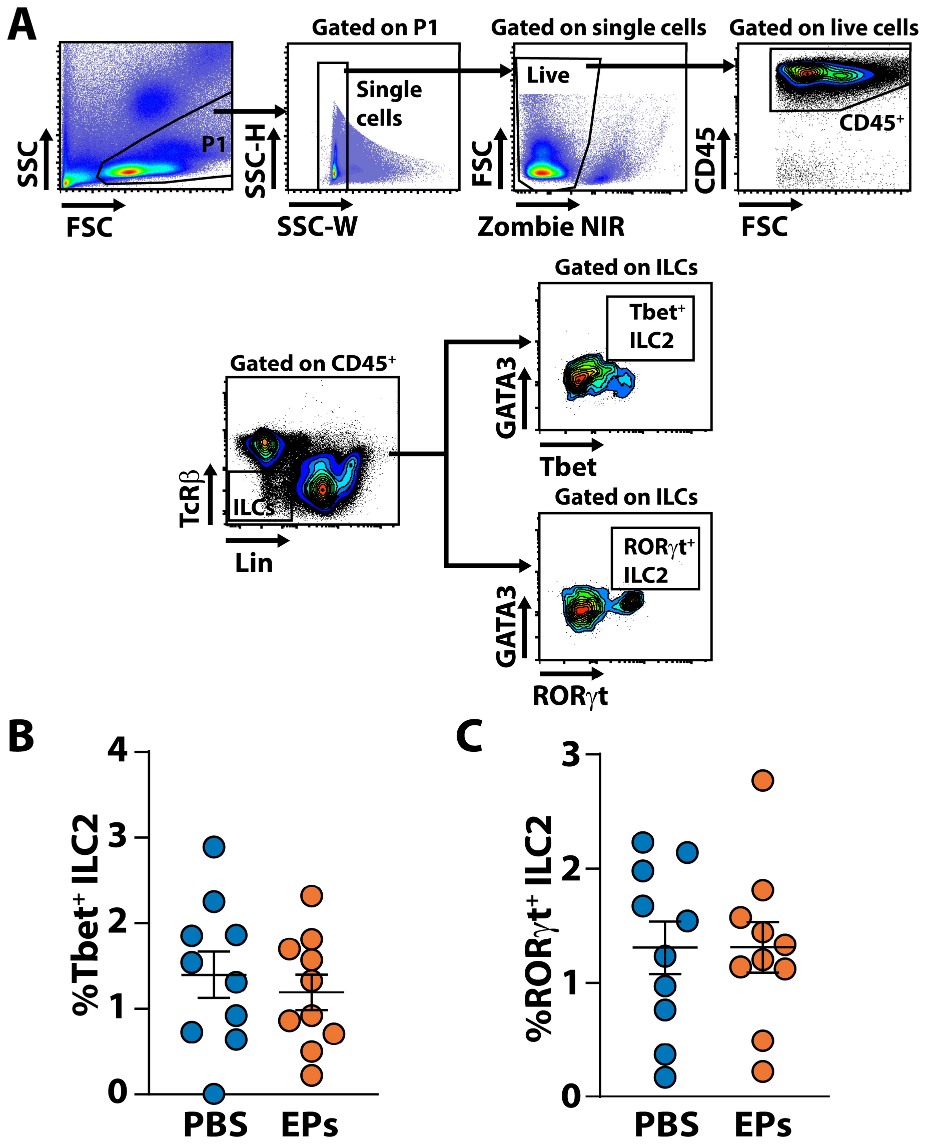


**Figure S8: Analyses of ILCs conversion.** **(A)** Gating strategy to identify Tbet$^{+}$ ILC2 and ROR$\gamma$t$^{+}$ ILC2 and quantification of **(B)** Tbet$^{+}$ ILC2 and **(C)** ROR$\gamma$t$^{+}$ ILC2 in the lungs of mice intratracheally instillated with 10 µg EPs in 50 µL PBS (n=10) or with 50 µL PBS alone (n=10) for 7 days. Data are presented as scatter plots with mean±SEM. Differences between two groups were evaluated by using the Wilcoxon-Mann–Whitney test.

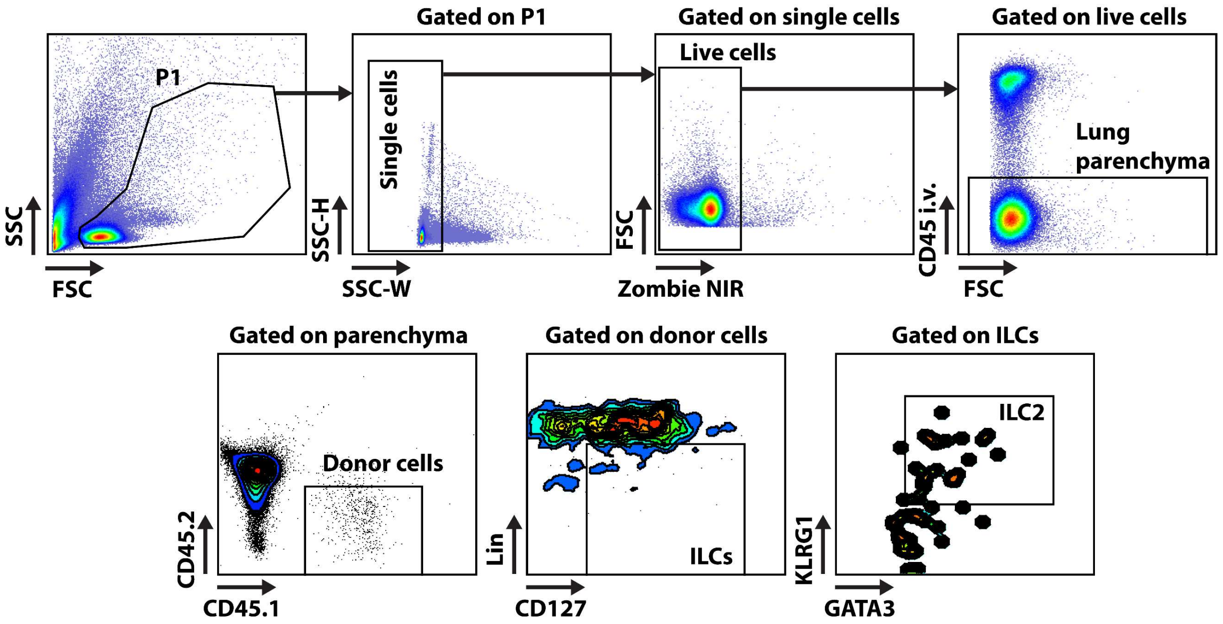


**Figure S9: Flow cytometry gating strategy used for the identification of donor derived ILCs in the lungs of receiver mice.** Donor derived cells in the lung parenchyma were identified as negative for CD45 i.v., negative for CD45.2 and positive for CD45.1. Then ILC2 were identified as previously described.

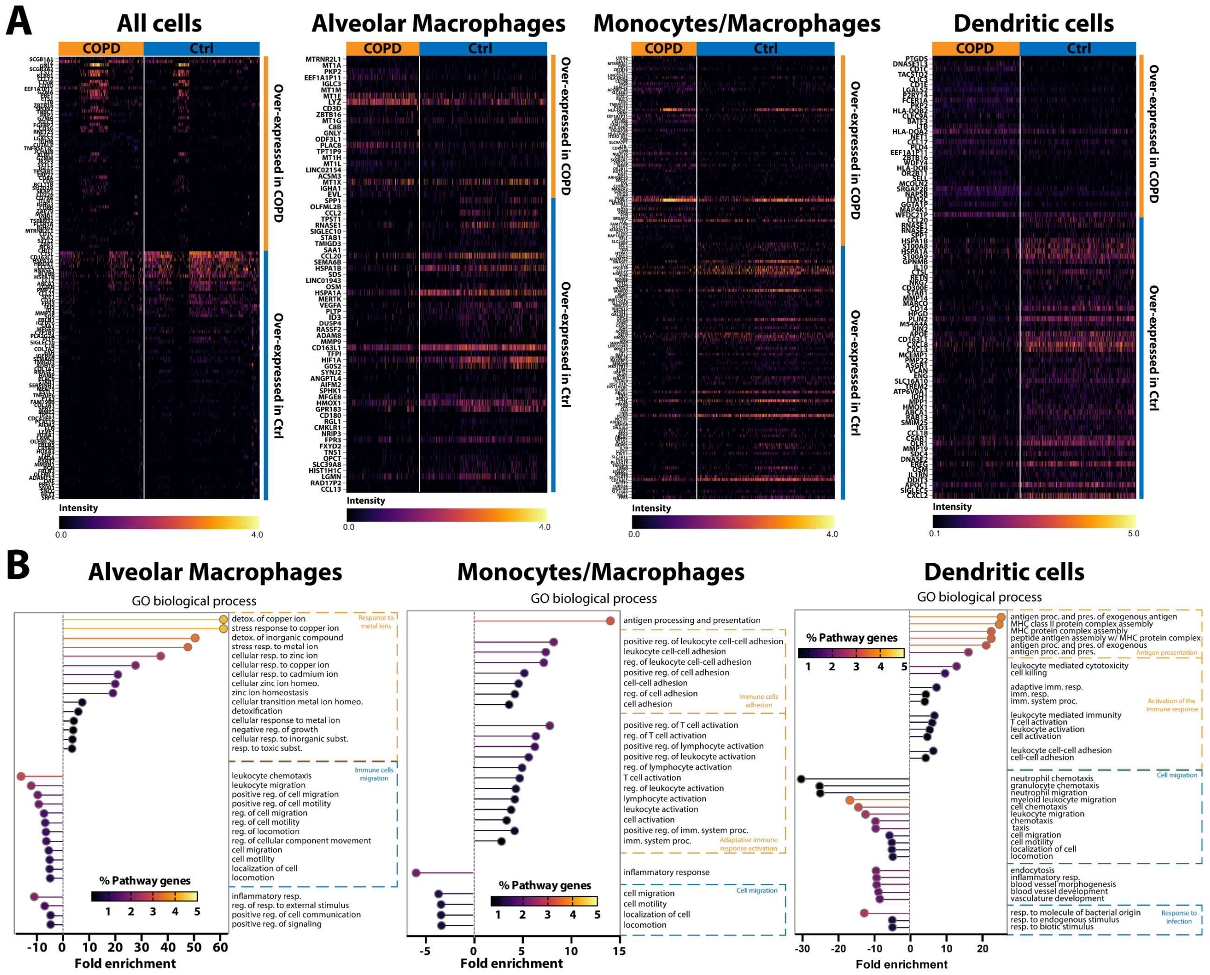


**Figure S10: Supplemental scRNAseq analyses. (A)** Heatmaps of differentially expressed genes between patients with COPD and healthy donors in all cells, alveolar macrophages, monocytes/macrophages and dendritic cells. **(B)** Gene ontology analysis of pathways up- or down-regulated in alveolar macrophages, monocytes/macrophages and dendritic cells.

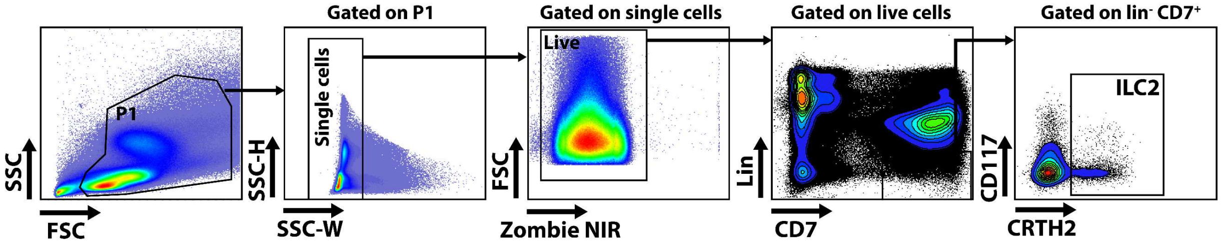


**Figure S11: Flow cytometry gating strategy used for the identification of ILC2 in patients.** ILC2 were identified as $Lin^{-}CD7^{+}CD117^{+}CRTH2^{+}$.

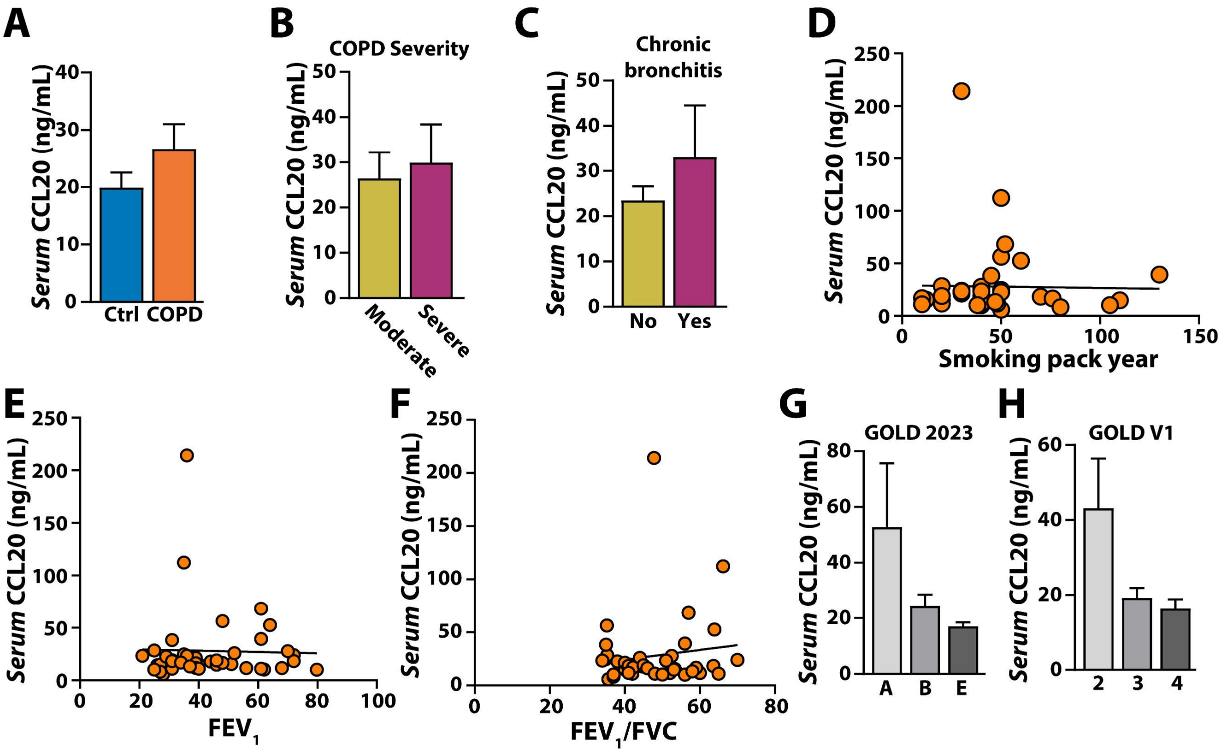


**Figure S12: Supplemental analysis of CCL20 levels in the serum of patients with COPD. (A)** CCL20 serum level in healthy donors and patients with COPD. **(B)** CCL20 serum level in patients with moderate or severe COPD. **(C)** CCL20 serum level in patients with COPD with or without chronic bronchitis. **(D)** CCL20 serum level in patients with COPD according to smoking history. **(E)** CCL20 serum level in patients with COPD according to $FEV_1$. **(F)** CCL20 serum level in patients with COPD according to $FEV_1/FVC$. **(G)** CCL20 serum level in patients according to GOLD 2023 COPD ranking. **(H)** CCL20 serum level in patients according to GOLD V1 COPD ranking.

| | COPD n=41 | Controls n=13 |
|---|---|---|
| Demographics | | |
| Sex, n (male/female) | 19/22 | 9/4 |
| Age (years) | 62.5 ± 7.2 | 58.3 ± 6.1 |
| BMI, kg/m² | 25.1 ± 6.8 | NA |
| Smoking history | | |
| Never smokers | 0% (0)* | 61.5% (8) |
| Ex-smokers | 70.7% (29)* | 15.4% (2) |
| Smokers | 29.3% (12) | 23.1% (3) |
| Pack-years | 45.4 ± 25.3 | NA |
| Inhaled treatment | | |
| LAMA and/or LABA | 41.46% | NA |
| ICS + LAMA and/or LABA | 51.22 | |
| Pulmonary function tests | | |
| FEV1, % | 43.99 | NA |
| FVC, % | 90.28 | |
| TLC, % | 132.28 | |
| RV, % | 226.26 | |
| Spirometric GOLD 1/2/3/4 | 0/13/19/9 | |
| GOLD A/B/E | 7/17/16 | |

**Table S1: Characteristics of the study population.** Values are expressed as mean ± Standard Error of the Mean (SEM) or percentage (number) **BMI**: Body Mass Index, **GOLD**: Global Initiative for Chronic Obstructive Lung Disease, **LABA**: long-acting beat-agonist, **LAMA**: long-acting muscarinic antagonist, **NA**: Not available, **ICS**: inhaled corticosteroid, **FEV1**: forced expiratory volume in one second, **FVC**: forced vital capacity, **TLC**: total lung capacity, **RV**: residual volume. *$p < 0.05$ vs control group.
